# Comprehension precedes production of a complex call sequence

**DOI:** 10.64898/2026.06.22.733898

**Authors:** Stephanie L. Mason, Amanda R. Ridley

## Abstract

Growing evidence of animals combining discrete, meaningful calls into sequences—a feature once thought unique to linguistic syntax—has presented the opportunity to investigate the evolutionary origins of syntactic communication. The arbitrary assignment of meaning to words marks an important step in human language evolution, and a necessary precursor to generating further meaning through sentences. Studying how other animals that produce call sequences learn the meaning of these signals could help shed light on how referentiality and semantic combinatoriality evolved. Given the presence of meaningful call sequences has only recently been revealed in several non-human animals, ontogenetic studies of the comprehension of these vocalisations are, to date, non-existent. Western Australian magpies (*Gymnorhina tibicen dorsalis*) combine discrete calls into a diverse array of call sequences. Recent evidence shows these sequences are socially learned, but the developmental stage at which fledglings respond correctly to them remains unstudied. We performed playbacks of a discrete alarm call and call sequence to fledglings over the course of their first 18 weeks out of the nest, identifying when they differentiate between the low-level disturbance associated with the discrete call and the high-grade aerial threat associated with the sequence. Fledglings showed immediate vigilance to both vocalisations but exhibited significantly greater vigilance and upward scanning following the sequence. Critically, fledglings showed this response to the sequence from the first week of testing, with no effect of age on the response to either vocalisation. These findings suggest that comprehension precedes production of sequences in magpies and that sequence meanings are either learned rapidly or have an innate basis. While further investigation is essential, this study offers the first empirical insight into the ontogenetic emergence of combinatorial comprehension in a non-human animal.

## Introduction

The combining of sounds into meaningful units and those units into structured sequences, termed *syntax*, was long thought absent outside of human language (Hurford, 2012; Suzuki et al., 2020). As a result, the evolutionary origins of this crucial feature of language remain unresolved (Christiansen & Kirby, 2003; Fischer & Price, 2017; Hauser & Fitch, 2003; Weiss & Newport, 2006). However, recent evidence in a wide array of non-human animals (hereafter *animals*) has revealed the presence of similar mechanisms of combining meaningful calls—sometimes comprised of smaller vocal elements (Walsh et al., 2023)—into sequences (Berthet et al., 2019, 2025; Bosshard et al., 2024; Collier et al., 2020; Déaux et al., 2016; Engesser et al., 2016; Girard-Buttoz et al., 2022; Hedwig & Kohlberg, 2024; Ouattara et al., 2009; Suzuki et al., 2016). This growing body of research presents the opportunity to investigate the potential evolutionary origins of syntactic communication through comparative research across taxa. However, while there is now substantial evidence for the presence of semantic combinatoriality, there is still little known about how this feature first emerges in the species in which it is seen.

To date, research on the emergence of meaning comprehension is limited to non-combinatorial signals, with a particular focus on alarm calls (Davis, 1984; Deshpande et al., 2022; Hollén & Manser, 2006; León et al., 2023; Seyfarth & Cheney, 1980; Wilson & Hare, 2006). Given that responding correctly to alarm calls can be crucial for survival, many species exhibit a degree of innate response to these vocalisations (Hollén & Manser, 2006; Lind & Cresswell, 2005; Seyfarth & Cheney, 1980, 1986). In social species, this is generally seen in the form of innate but indiscriminate vigilance to alarm calls before more specific, adult-like responses are learned from observing the behavioural responses of social contacts (Hollén & Manser, 2006; Seyfarth & Cheney, 1980, 1986; though see Göth, 2001, 2002 for evidence of innate predator-specific responses in a precocial species)—a process termed vocal comprehension learning (Janik & Slater, 2000). Even when social learning opportunities are available, the high cost of relying on a trial-and-error approach up to the point of learning still appears to favour some degree of innate vigilance to alarm calls (Lind & Cresswell, 2005). Whether this pattern extends to more complex, combinatorial threat vocalisations, however, remains unstudied. By comparing how comprehension of individual signals and the sequences in which they are combined emerges during ontogeny, we can explore whether combinatoriality arises from distinct cognitive mechanisms or builds on those involved in learning the meaning of discrete calls. This would provide crucial insight into how complex combinatoriality may have evolved from simpler referential signals.

Most research into combinatoriality has focussed on a small number of vocalisations of two sounds or signals at a single structural level—either the combination of acoustic elements within a call (e.g., chestnut-crowned babblers, *Pomatostomus ruficeps*, Engesser et al., 2019) or between calls in two-call combinations (e.g., the alert-recruit combinations of southern pied babblers, *Turdoides bicolor*, Engesser et al., 2016; and Japanese tits, *Parus minor*, Suzuki et al., 2016, but see also Collier et al., 2020; Déaux et al., 2016; Hedwig & Kohlberg, 2024; Ouattara et al., 2009). While such studies have been invaluable in revealing the extent of combinatoriality across the animal kingdom, it remained unclear whether further structural complexity and repertoire flexibility more akin to the generative nature of linguistic syntax may be present in animal vocal systems. By employing whole-repertoire approaches, several recent studies have revealed extensive repertoires of over 100 unique sequences, with some sequences exceeding 10 calls, providing the opportunity for potentially closer parallels to be drawn with linguistic syntax. Specifically, extensive repertoires of long, complex sequences—often nine or more calls—have been discovered in common marmosets *Callithrix jacchus* (Bosshard et al., 2024), chimpanzees *Pan troglodytes* (Girard-Buttoz et al., 2022), and Western Australian magpies (Walsh et al., 2023), with the latter being the only species to date shown to use predictable ordering rules to combine both sounds within calls and calls within sequences (Mason, Walsh et al., 2026; Walsh et al., 2023; Figure 1). While evidence of call sequences in two distinct primate groups provides direct insight into the emergence of combinatoriality within the primate lineage—likely evolving some 45 million years ago (Leroux & Townsend, 2020)—evidence in much more distantly related groups like songbirds provides the opportunity to identify the drivers of this capacity independent of shared ancestry (Suzuki, 2021). Additionally, magpies are one of very few species that, alongside combinatoriality, also possess the capacity for open-ended vocal production learning—the ability to learn to produce new vocalisations from heard models throughout life (Kaplan, 1999; Suthers et al., 2011)—a feature crucial to the complexity of human language (Verpooten, 2021). Given this capacity is seemingly absent in non-human primates (Fischer et al., 2015; Janik & Knörnschild, 2021), this places magpies as a powerful comparative tool for investigating the origins of complex combinatoriality in a species that, like us, is theoretically unconstrained by repertoire size.

**Figure 1.**
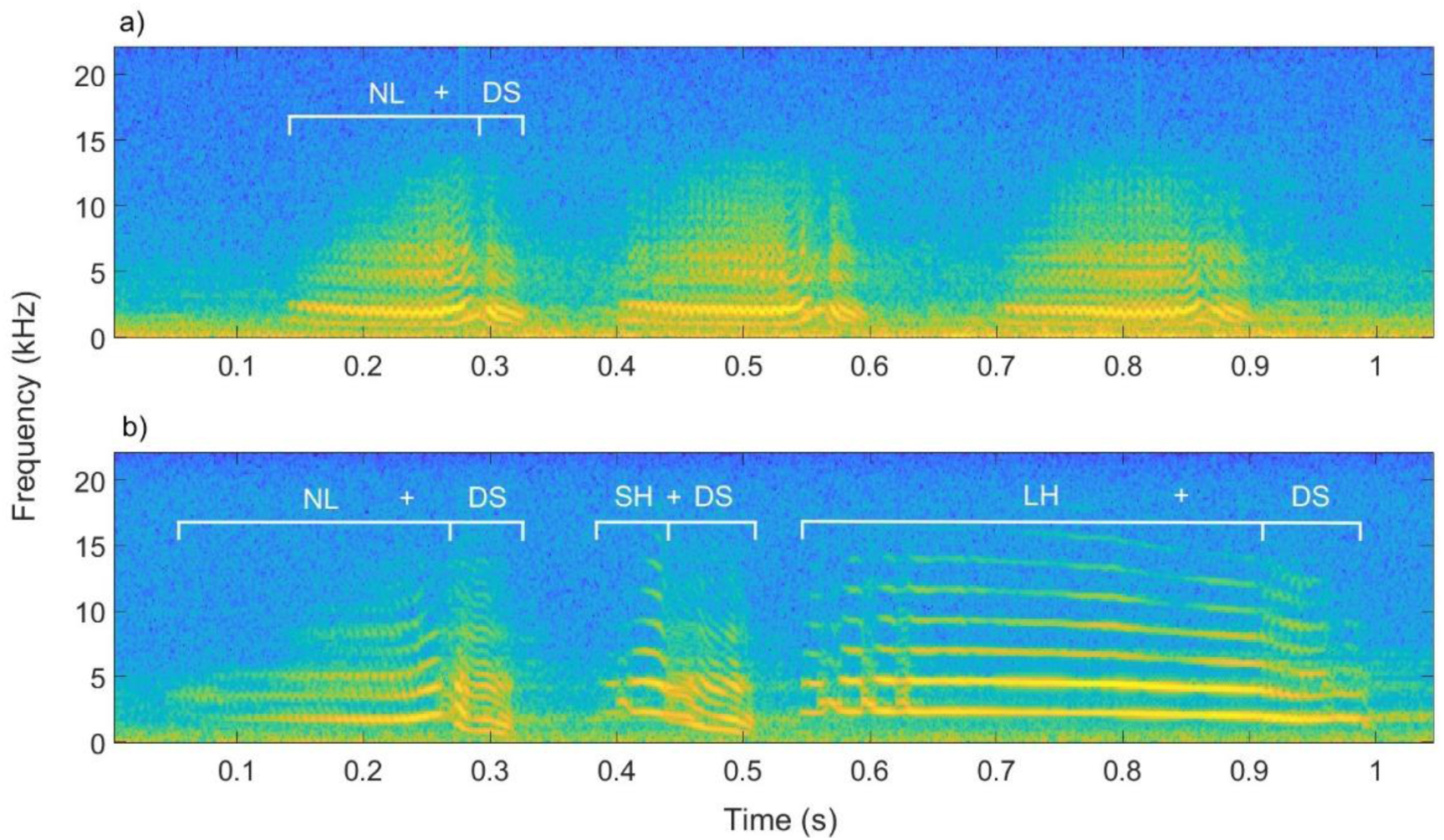
Examples of magpie vocalisations with (a) showing a series of discrete *NLDS* calls, and (b) showing an *NLDS-SHDS-LHDS* sequence. Vocalisations shown were recorded at 44.1kHz sample rate and spectrograms were generated with a 256-point FFT with a Hann window and 250-sample overlap leading to a frequency resolution of ∼172.3Hz. Segments (the smallest vocal unit) are combined into discrete calls such that segments within the call are distinguishable only by sudden spectral shifts (e.g., *NL + DS* in *NLDS*). Discrete calls can either be produced in isolation or in series such as the ‘*NLDS*’s in (a)—such that they are separated from repeats by periods of silence ≤0.5s—or combined into call sequences where individual calls are separated by periods of silence ≤0.5s. Individual vocal events are those that occur >0.5s apart. Segments are labelled according to spectrographic structure confirmed using spectrographic cluster analysis; refer to Walsh et al., 2023 for further detail on magpie vocal classification.

A recent ontogenetic study of magpie fledglings revealed that their call sequence repertoire is socially learned, with fledglings producing the sequence repertoire of their social group, not the broader population, and more sociable fledglings producing more sequences, earlier in development (Mason, King & Ridley, 2026). In contrast, the much smaller number of discrete calls that form these many sequences (five discrete calls versus 119 call sequences recorded across the population) are produced as early as the first week post-fledging (eight weeks earlier than the average onset of sequences), with little variation in discrete call repertoire between groups or fledglings, and no effect of sociability on their development (Mason, King & Ridley, 2026). While it is possible that discrete calls are simply learned during the nestling phase or even earlier (see Kleindorfer et al., 2024 for example of in-ovo song learning), their early onset, paired with there being little variation in discrete call repertoire between individuals and no effect of social interaction on call development, suggests a possible innate basis to discrete magpie calls (see also Mason, 2025 for acoustic analysis supporting their innate basis). Having shown that calls and sequences emerge at different stages of development and seemingly through different cognitive processes (i.e., innate versus learned), investigating how fledglings come to understand the function of each would provide valuable insight into the ontogenetic foundations of combinatoriality.

Here, we performed playbacks of a discrete call associated with low-grade general disturbance (Dutour et al., 2020, 2023; Walsh, 2024) and a call sequence that appears to be associated with high-grade aerial threat contexts (Walsh, 2024) to fledgling magpies over the course of development. Our aim was to ascertain (i) when fledglings first respond, regardless of how, to these vocalisations and (ii) when fledglings first show differentiated responses to these vocalisations—i.e., scanning the sky for aerial threats following the sequence but not the discrete call. We predicted that fledglings would initially show innate, indiscriminate vigilance to both vocalisations. As the fledglings aged, however, their responses to the vocalisations would diverge, as they began to show increased upward scanning following the call sequence as they learned from observing group members, its association with aerial threats.

## Methods

### Study population

Western Australian magpies are sexually dimorphic passerines that live in stable, cooperatively breeding social groups (Pike et al., 2019). Magpies produce complex vocalisations from synchronised song (Dutour & Ridley, 2020) and mimicry (Kaplan, 1999; Suthers et al., 2011), to discrete calls and call sequences (Dutour et al., 2020; 2023; Walsh et al., 2019, 2023; 2024). Hatchling magpies spend 3-4 weeks in the nest being provisioned before fledging (Pike et al., 2019). The post-fledging dependent period represents an extended period of offspring development where young continue to beg for and be provisioned with food by adult group members (Ashton et al., 2018; Edwards et al., 2015). Due to high fledgling mortality rates, a limited number of fledglings were available for playback experiments. Playbacks were conducted on 9 fledglings from 7 wild magpie groups situated in urban areas in the Perth metropolitan suburbs of Crawley (31.97°S, 115.82°E; n = 4 groups) and Guildford (31.89°S, 115.97°E; n = 3 groups) all of whom were concurrently being observed and recorded as part of a larger ontogenetic study (Mason, King & Ridley, 2026). The groups are part of a study population used for long term research (founded by ARR) and are habituated to human presence, allowing for natural observation and recordings to take place from less than 10m (Speechley et al., 2024; Walsh et al., 2023). Individual birds are identifiable through unique coloured rings or physical features (scarring or plumage).

### Call collection and preparation

Magpies produce 4 distinct vocal ‘segments’—noisy line (*NL*), short high (*SH*), long high (*LH*) and down sweep (*DS*). At the first level of structure, these segments are combined in a process akin to phonological syntax to form discrete calls where constituent segments are delineated only by sudden spectral shifts and combined using predictable first-order rules—where each segment depends on the one prior (Walsh et al., 2023; Figure 1a). At the next structural level, discrete calls are combined into call sequences using at least first-order rules (Walsh et al., 2023; though see Mason, Walsh et al., 2026 for recent evidence of second-order structure) where individual calls are separated by brief periods of silence ≤0.5s (Walsh et al., 2023; Figure 1b). The vocalisations used as stimuli in the current study were the ‘noisy line-down sweep’ (*NLDS*) call formed from two segments (*NL* and *DS*) and a call sequence containing the same discrete call along with two others: ‘noisy line-down sweep + short high-down sweep + long high-down sweep’ (*NLDS-SHDS-LHDS*) (Walsh et al., 2023; Figure 1).

*NLDS* is well established as a low-grade general disturbance alarm call (Dutour et al., 2020; 2023; Walsh, 2024) used in contexts such as the approach of domestic dogs or people, both of which magpies can easily evade. At the start of the current study, concurrent research into the function of magpie sequences was still ongoing but *NLDS-SHDS-LHDS* was thought to be associated with high-grade aerial threats (personal observation; correspondence with S. L. Walsh) with support for this function shown in adult males in the concurrent research (Walsh, 2024), following completion of our study. Females, however, were found to predominantly produce *NLDS-SHDS-LHDS* in seemingly nil threat contexts and during recruitment of group members (Walsh, 2024). Given the absence of a threat cannot be confirmed by observers limited to identifying threats visible from the ground (magpies have the vantage point of flight and perching high in the tree canopy), and that recruitment is common during high-grade aerial threats (both conspecific intruders and the presence of predatory birds), the sex difference observed may simply be a result of observers failing to identify an aerial threat that was present. Nevertheless, we quantify time spent scanning upward versus horizontally as a means to further test the aerial threat association of this sequence in fledglings and directly test for differences in fledgling responses to male versus female productions of the sequence to address potential sex differences in function. Sample recordings of *NLDS* (hereafter ‘the alarm’) and *NLDS-SHDS-LHDS* (hereafter ‘the sequence’) were taken from opportunistic recordings of group members prior to the start of the study or from existing recordings of still-living members of the study groups held in a sound database collected between 2019-2022 by SLM and S. L. Walsh. All recordings were made using a RØDE NTG-2 directional microphone in a Blimp suspension windshield system, and a Roland R-05 or R-07 wav/MP3 recorder set at a sampling rate of 44.1 kHz.

Using Adobe Audition 24.4.1.3, individual productions of the alarm and sequence with good signal to noise ratio (determined visually based on clear signal on spectrogram) were identified and extracted for use in playback stimuli. One caller was selected from each of the tested fledglings’ groups based on the number of available high-quality samples from that individual. For some groups, the identity of the caller for the alarm and sequence treatments was the same, while for others it was only possible to keep the identity consistent within each treatment. There were two groups where two fledglings were followed to boost sample size, but in both instances the fledglings were from different mothers and were always tested in isolation (i.e., geographically isolated within the group territory and at different times). At one of these groups, the same set of alarm call stimuli produced by an adult male had to be used for both fledglings, but a different set of sequence stimuli were available for each fledgling, from two different adult females. At the other group, the same stimuli were used for both treatments for both fledglings; however, the fledglings were rarely in contact with one another due to the large size of the group, and were different ages so were never tested in the same week. For one fledgling, there were no sequence recordings available, so only the alarm and control treatments could be conducted. Due to limited high-quality recordings available, it was not possible to keep the sex of the caller consistent across all groups, so sex of caller for each set of stimuli was later evaluated to test for differences in fledgling responses to male versus female callers. None of the stimuli contained vocalisations of the mother of the fledgling being tested, in case this produced a different response than to other group members. The influence of the father was not considered as extra-group paternities are high in this species (> 85 %; Hughes et al., 2003) and thus the father is often not a group member or known to fledglings. In contrast, mothers are readily identified through nest-building and incubation behaviour. In some instances, where recordings of the sequence and alarm were limited, the vocalisations within the stimuli were artificially formed from combining individual productions of *NLDS* and *SHDS-LHDS* (a shorter sequence that is produced in isolation as well as within the target sequence, *NLDS-SHDS-LHDS)* from the same caller, with inter-call distance reflecting the mean duration for periods of silence observed in natural productions of the sequence (0.1 seconds; Walsh et al. 2023) (N = 11 of the sequence stimuli), or by cutting the *NLDS* from the start of a sequence in the case of the alarm stimuli (N = 3 of the alarm call stimuli). Previous work has revealed discrete productions of *NLDS* to be indistinct from *NLDS* produced within sequences (Walsh et al., 2019), and vigilance to artificially created sequences (using the same method as the present study) to be no different than to natural productions of the same sequence (Walsh, 2024). However, natural versus artificial sequences were still compared during analysis to assess whether this impacted response. For most fledglings, the final number of stimuli available was four per experimental treatment (x̄ = 3.8 alarm call stimuli, range = 3 to 4 across 9 fledglings, x̄ = 3.4 sequence stimuli, range, 2-4 across 8 fledglings).

A high-pass filter was used on all stimuli to filter out background noise <0.3kHz to improve signal quality and avoid excessive low-frequency noise in conjunction with the natural soundscape during playback. The range of maximum amplitude for productions of the alarm and the sequence by adult magpies measured at ∼1m from the caller is between 68dB-73dB SPL (A-weighted, re 20 µPa, measured using a DIGITECH QM-1591 sound level meter; Walsh, 2024). Given that higher amplitudes may elicit stronger responses (Walsh, 2024), and the aim of this study was to test for differences in behavioural responses to the meaning of each vocalisation, maximum amplitude was kept consistent across both treatments. As such, each recording used was normalised so that when played back at maximum volume on the speaker, maximum amplitude was 70dB SPL (A-weighted, re 20 µPa, measured ∼1m from the speaker).

From observations made in-field, adults will often ignore a single alarm call and only show an observable reaction when multiple productions of the same call are given. Previous work has shown magpies respond to urgency information contained within repetitions (Dutour et al., 2023). While this was ascertained from repeating calls within the vocalisation/bout (i.e., *NLDS-NLDS-NLDS-NLDS* where each call is separated by <0.5s, Dutour et al. 2023), repeated distinct vocalisations (separated by >0.5s) in quick succession likely have a similar effect. As such, each stimulus contained the same vocal sample repeated three times within the audio track, with each repeat separated by a ∼2 second interval (within the range of natural intervals observed between repeat productions of the same vocalisation from the same caller; personal observation) of low-level background noise containing no other magpie or heterospecific calls, taken from the same source recording as the sampled vocal. This allowed for a large enough behavioural response to occur such that variation between playbacks would be discernible, and meant relative urgency was kept consistent across treatments. Within treatments, all stimuli were the same length (to the nearest second) from the start of the first, to the end of the third repeat.

### Control playback treatment

A contact call of the Australian wood duck (*Chenonetta jubata*) was used as a control to ensure the playback set up itself was not the cause of any response. This species is naturally present at all groups in the study population and does not produce a vigilance response from adult magpies (personal observation). Three sample recordings were collected opportunistically from ducks present at one of the Guildford magpie groups that form part of the study population but that was not being used in the present study (so that the control would be equally unfamiliar to all tested individuals should there be identity information encoded within wood duck calls that is recognisable to magpies). Three stimulus tracks were created, one for each of the sample recordings, with the sampled call repeated thrice within the track, as with the experimental treatments. Playing the control back at the same amplitude as the experimental treatments was not possible as this would have been unrealistically loud for a natural duck call and may have produced an unnatural vigilance response as a result. Instead, all 3 control stimulus tracks were normalized to 20dB SPL (A-weighted, re 20 µPa, measured at ∼1m from the speaker) (within the range of natural productions of wood duck calls; personal observation).

### Data collection

Playbacks were conducted from the beginning of the breeding season in August 2022, every 3 weeks from 3 to 18 weeks post-fledge. Three weeks post-fledge was the earliest that fledglings were isolated from caregivers for sufficient time to conduct the playbacks. Only one fledgling was successfully tested after week 18 (due to fledgling mortality and/or group disturbances during visits) so only playbacks up to week 18 were included in the final dataset. On each of the test weeks, playbacks of both experimental treatments and the control treatment were conducted with each fledgling, with a minimum of 20 minutes between the end of the behavioural response or end time of the last playback (whichever finished last) and commencing playback of the next playback treatment. This allowed for normal behaviour to resume; however, order of playback treatment was still evaluated to assess if there was a cumulative effect of consecutive treatments on fledgling response on each test day (i.e. if they become habituated and less responsive, or indeed more physiologically aroused and more responsive, with each consecutive treatment). The stimulus track used for each test and the order in which the treatments were tested were both counter-balanced across each week of testing to avoid habituation to stimuli and cumulative response affecting results.

To test the impact of sociability on playback response, social interaction data from a concurrent observational study (Mason, King & Ridley, 2026) was used. This data was collected by recording all contacts within 10m of the fledglings throughout the course of hour-long focal follows that were conducted weekly from week 1 to 15 post-fledging, and then every 3 weeks until week 30 post-fledging (Mason, King & Ridley, 2026). Only the social data collected up until the final playback for each fledging was included in the analysis for the current study, to avoid skewing the results with data not reflective of their sociability at the time at which they were tested.

### Playback protocol

A portable speaker (Ultimate Ears BOOM 2) was used and placed on the ground 10m from the test individual concealed by leaf litter or topography such that it was not visible to the test individual. The speaker was positioned such that the playback would come from the direction of the calling individual heard in the stimulus to avoid a violation of expectation (i.e. where the playback is coming from a different direction than the location of the actual bird at the time of playback). The placement of the speaker on the ground also meant that upward scanning by the fledglings was a clear indication of them searching for aerial stimuli, and not just them looking to the source of the sound. Playbacks were only conducted once all group members were more than 10m from the fledgling (considered minimum distance for social isolation: Speechley et al., 2024) and the speaker, and the fledgling was foraging and not vigilant. Playbacks were only conducted when no major disturbance (e.g., presence of aerial predator) had occurred for at least 30 minutes, and no group members had produced any alarm/alert calls or sequences for at least 5 minutes. A handheld Panasonic HC-V520M video camera was used to film playback trials and recording was started at least 2 minutes prior to the playback to verify that the fledgling was non-vigilant. When at least 2 minutes of recording had been conducted and the fledgling was actively foraging with its bill down (clear sign of non-vigilance), the stimulus was played from an iPhone 11 connected to the speaker via an AUX cable to avoid signal compression or degradation. During experimentation the researcher, SLM, stood 10m from the speaker and the fledgling, only moving if necessary to maintain visual contact with the fledgling following commencement of the playback. Video recording continued until 2 minutes following the playback or until fledgling non-vigilance resumed if it had not done so by 2 minutes following playback. A total of 130 trials were conducted (alarm n = 44, sequence n = 41, control n = 45).

### Ethical note

Our data collection protocol was reviewed and approved by the University of Western Australia Animal Ethics Committee (Protocol number: RA/3/100/1656 and 2021/ET000272). All ringed birds had been previously ringed as part of the broader research project and not specifically for the current study. A maximum of two playbacks (each involving three treatments) were conducted in the same week at any group (five of the seven groups only experienced one playback per week) with a minimum gap of 20 minutes between treatments to reduce stress to the birds. Fledgling isolation from group members by ≥10m was achieved by waiting for this to occur naturally and at no point were birds disturbed to cause isolation.

### Response variables

During scoring of responses, all videos were muted and left unnamed, so it was not clear which treatment was being viewed or when playback had started. SLM then marked the start and end of periods of vigilance using Adobe Premier Pro 23.2.0, along with durations of time spent scanning horizontally versus upwards, and the start and end times of any fleeing or mobbing behaviour. Normal behaviour was considered resumed once the fledgling had spent at least 3 seconds foraging following vigilance, even if they then became vigilant again. This was because it could not be discerned if vigilance following a bout of foraging was due to the playback or other environmental variables. Following the blind appraisal of videos, the start and end of the playback was marked in each video for calculating latencies and durations of vigilance since playback start. We intended to score and analyse vocal response, as well as approach to speaker, fleeing from speaker, joining mobbing response and time spent vigilant. However, there were very few instances of vocal responses (i.e., any vocal type emitted by the fledgling following playback; n = 10 of 86 experimental treatments conducted), approach to speaker (n = 0), fleeing (n = 17), or mobbing (n = 11), so it was not feasible to analyse these responses. As a result, only (i) latency to react (i.e., how quickly they react), (ii) time spent vigilant (i.e., how long they react from the start of playback) and (iii) proportion of vigilance spent scanning up (versus horizontally, as an indication of scanning for an aerial predator as opposed to general disturbances) were scored and later modelled as response terms. In cases where normal behaviour did not resume within 2 minutes of the playback, visual contact with the fledgling was often lost, making it unclear how long the vigilance response lasted. As such, the minimum of 2 minutes of recording following playback was used as the cut off so that maximum latency to, and duration of vigilance possible, was 120 seconds. To assess repeatability of measurements, a second observer scored a random subset of 25 videos (20%). This was compared to SLMs scoring using the R package rptR (Nakagawa & Schielzeth, 2010) which showed very high repeatability for latency to react (R = 0.897 ± 0.05; 95% CI = 0.77, 0.95), time spent vigilant (R = 0.996 ± <0.01; 95% CI = 0.99, 1.00), and proportion of vigilance scanning up (R = 0.996 ± <0.01; 95% CI = 0.99, 1.00). Of the full dataset, one control trial was excluded because group members began alarming at a dog during the playback, and 2 control trials and 1 alarm trial were excluded because the fledgling became vigilant to passers-by as the playback started. The final dataset for the analysis of latency to react and time spent vigilant included 126 trials (n = 43 alarm trials, n = 41 sequence trials, n = 42 control trials). For some of these trials, time spent scanning up versus horizontally could not be ascertained as the fledgling’s head was obscured, or they fled during their vigilance response (n = 9 alarm trials and 9 sequence trials). The final dataset for scanning behaviour contained 108 trials (n = 34 alarm trials, n = 32 sequence trials and n = 42 control trials).

### Statistical analysis

All statistical analyses were conducted using generalised linear mixed models (GLMMs) using the package *glmmTMB* (Brooks et al., 2017) in R v.4.2.3 (R Core Team, 2023). Individual ID was included as a random effect in all models. Group ID and caller ID were not included because they were highly confounded with individual ID, explained less variance, and caused singularity issues when added alongside individual ID. All continuous predictors were centred around zero (Harrison et al., 2018).

Latency to react and time spent vigilant from start of playback were right skewed and overdispersed, so were both modelled using a negative binomial distribution (zero-inflated models did not improve fit so were not used) with a logarithm link function. For the latency to react model, one alarm trial was excluded from the data as an outlier (identified using *DHARMa*). Proportion of vigilance spent scanning upwards (relative to horizontal scanning) was modelled using a binomial GLMM with a logit link function, where the response variable was the number of seconds spent scanning up versus horizontally.

All models were initially run using the full data set including both experimental treatments and the control treatment. Predictors included: treatment, trial order, stimulus track used (stimulus ID) and age of fledgling at time of testing. In Mason, King & Ridley (2026), an effect of average adult association (the average of the proportion of time that fledglings spend with each adult social contact) and proportion spent in contact (the proportion of total observation time spent with others and not alone, i.e. 1 - time alone) on the rate of learning and the diversity of sequences produced by fledglings. As such, these two variables were also included as predictors.

After running models including all three treatments, additional models were run using a subset of the data that excluded the control treatment. This was necessary to look at the effect of caller sex and artificial versus natural stimuli (termed ‘stimulus type’) on fledgling responses, as both were rank deficient in the control treatment level (i.e. all control stimuli were natural recordings of a single sex). Time spent vigilant was the only one of the three response terms for which the model excluding the control treatment yielded additional results (the reduced scanning dataset did not have sufficient power to include further predictors while maintaining stable model fit, and there was little variation in latency to react outside of the control treatment), so latency to react and proportion scanning up are not presented as control-excluded models.

### Model comparison

Akaike’s Information Criterion corrected for small sample size (AICc) was used to select between candidate models by comparing them to each other and the basic model (containing only the intercept and the random term). The top model sets contained all models within 2 ΔAICc of the top model. For models with similar AICc, the simplest model was selected (i.e. fewest terms, *sensu* Harrison et al., 2018). All predictor terms included together in a model had a variance inflation factor (VIF) < 3. Following model selection, pairwise comparisons of categorical terms were run for the top models using the *emmeans* package where relevant (Lenth, 2023). For the top model set, the packages *DHARMa* (Hartig, 2022) and *performance* (Lüdecke et al., 2021) were used to check model assumptions.

## Results

### Control-inclusive models

For all three response variables—latency to react, time spent vigilant, and proportion scanning up—treatment was the best predictor. Fledglings reacted almost immediately to both the alarm (x̄ = 0.33 seconds ± 0.13) and the sequence (x̄ = 0.46 seconds ± 0.10)—with no significant difference between the two (Table S.4i)—but were significantly slower to react to the control (x̄ = 21.43 seconds ± 6.89) than either experimental treatment, with some not reacting to the control at all (Table 1i; Figure 2a; see Table S.3. for pairwise comparisons). Significant differences in time spent vigilant were found between all treatments, with fledglings spending the longest time vigilant in response to playback of the sequence (x̄ = 48.63 seconds ± 6.65), then the alarm (x̄ = 20.88 seconds ± 2.30), and finally the control (x̄ = 7.12 seconds ± 0.86) (Table 1ii; Figure 2b; see Table S.5. for pairwise comparisons). Fledglings spent significantly more time scanning up during vigilance responses to sequence playbacks than both discrete call and control playbacks (Table 1iii; Figure 2c; see Table S.9. for pairwise comparisons).

**Figure 2.**
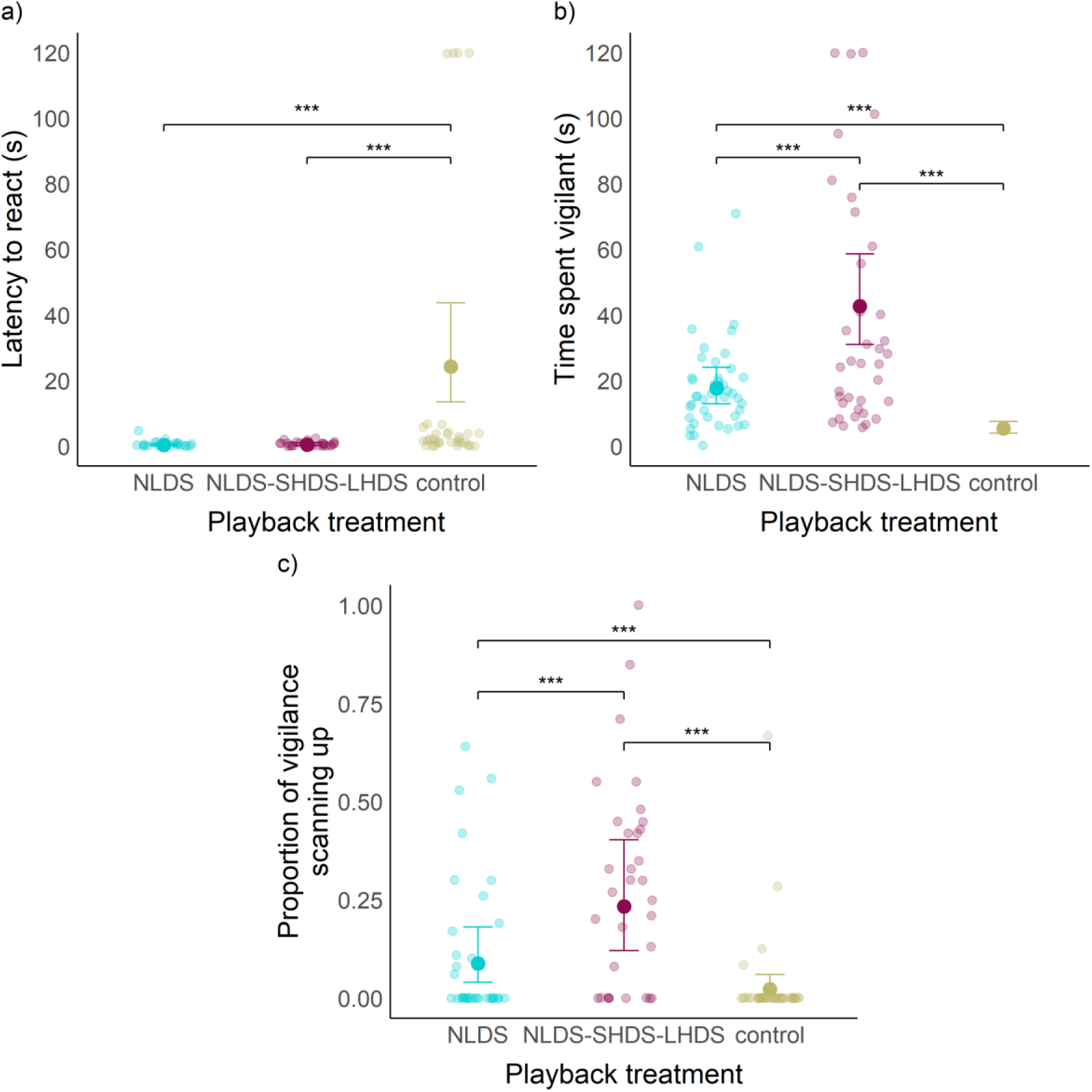
Relationship between playback treatment and (a) latency to react to (n = 125 trials), (b) time spent vigilant following the playback (n = 126 trials) and (c) proportion of vigilance spent scanning up (n = 108 trials). Treatment types: *NLDS* (a general disturbance alarm call), *NLDS-SHDS-LHDS* (a three-call sequence associated with aerial predator threats) and a control (an Australian wood duck contact call). Playbacks were conducted on 9 magpie fledglings from 7 groups when fledglings were between 3- and 18-weeks post-fledging. Pale dots depict raw data. Bars illustrate 95% confidence intervals (CI) for back-transformed estimated marginal means (shown by larger, opaque dots). Significance between treatments indicated by brackets with ‘*’ indicating p < 0.05; ‘**’ p < 0.01; and ‘***’ p < 0.001. Colours denote treatment level for raw data, CIs and estimated means.

**Table 1.**
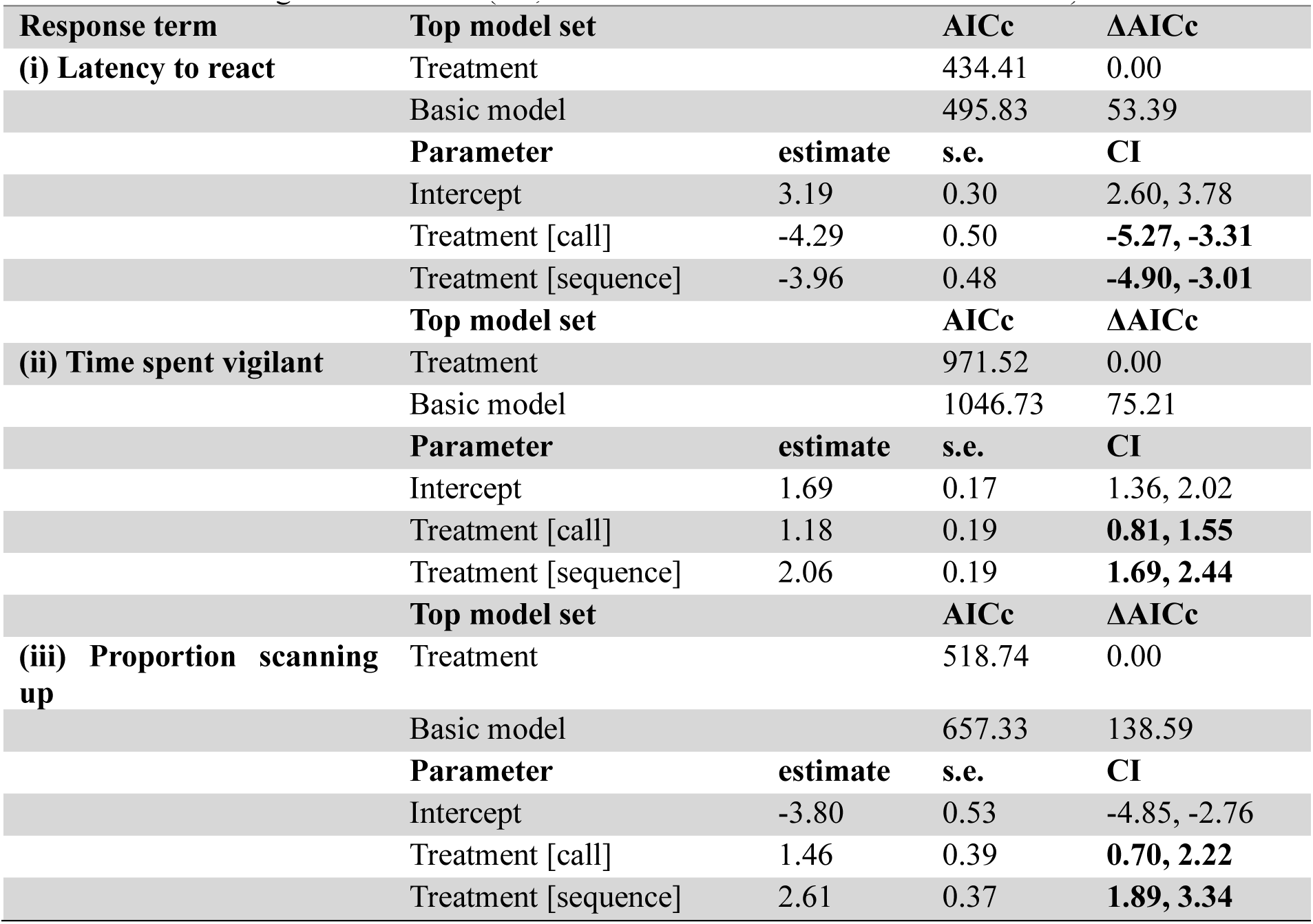
Top model set for fledgling (i) latency to react and (n = 125 trials), (ii) time spent vigilant (n = 126 trials) and (iii) proportion of vigilance spent scanning upwards following playbacks (n = 108 trials). Analyses were performed using negative binomial GLMMs for (i) and (ii) and a binomial GLMM for (iii). The reference level for treatment is control and fledgling ID was the random term for all models. Parameter coefficient estimates ± standard error (s.e.) and 95% confidence intervals (CI) are provided for each top model set; see electronic supplementary material for full model sets. Bold CI values indicate significant results (i.e., confidence intervals do not intersect zero).

### Time spent vigilant—control-excluded models

Looking at just the experimental treatments, alongside the greater time spent vigilant to the sequence than the alarm call (Figure 3a), caller sex affected time spent vigilant following playback: fledglings were vigilant for longer following both the alarms and the sequences of male callers than those of female callers (Table 2; Figure 3b). Order of playback also had a significant impact (Table 2), however, pairwise comparisons showed only the 1^st^ and 2^nd^, but no other trials, to be significantly different from one another—with longer vigilance following trials performed 1^st^ than 2^nd^ (supplementary Table S.6.). The minimal influence of trial order was further confirmed upon examining the means of both treatment and caller sex when averaged over the levels of trial order—vigilance remained significantly lower for alarm trials and for trials using female callers regardless of the level of trial order (supplementary Table S.5.; Figure S.1.).

**Figure 3.**
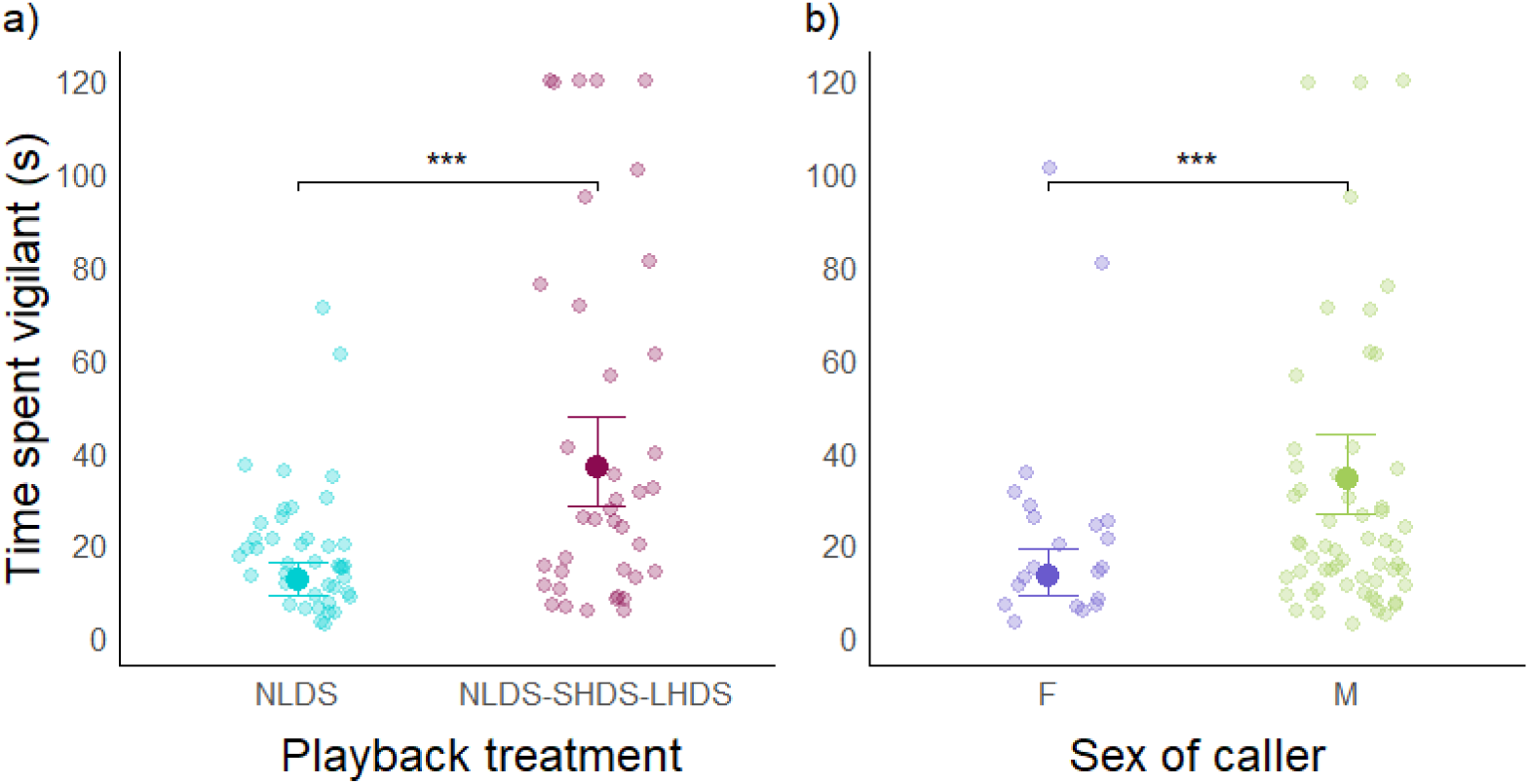
Effect of (a) treatment and (b) caller sex on time fledglings spent vigilant to playback trials (n = 84 trials). Treatment types included in (a) are *NLDS* (a general disturbance alarm call) and *NLDS-SHDS-LHDS* (a three-call sequence associated with aerial predator threats). (b) shows sex of the caller in the playback exemplar. Pale dots depict raw data. Bars illustrate 95% confidence intervals (CI) for back-transformed estimated marginal means (shown by larger, opaque dots). Significance between treatments indicated by brackets with ‘*’ indicating p < 0.05; ‘**’ p < 0.01; and ‘***’ p < 0.001.

**Table 2.**
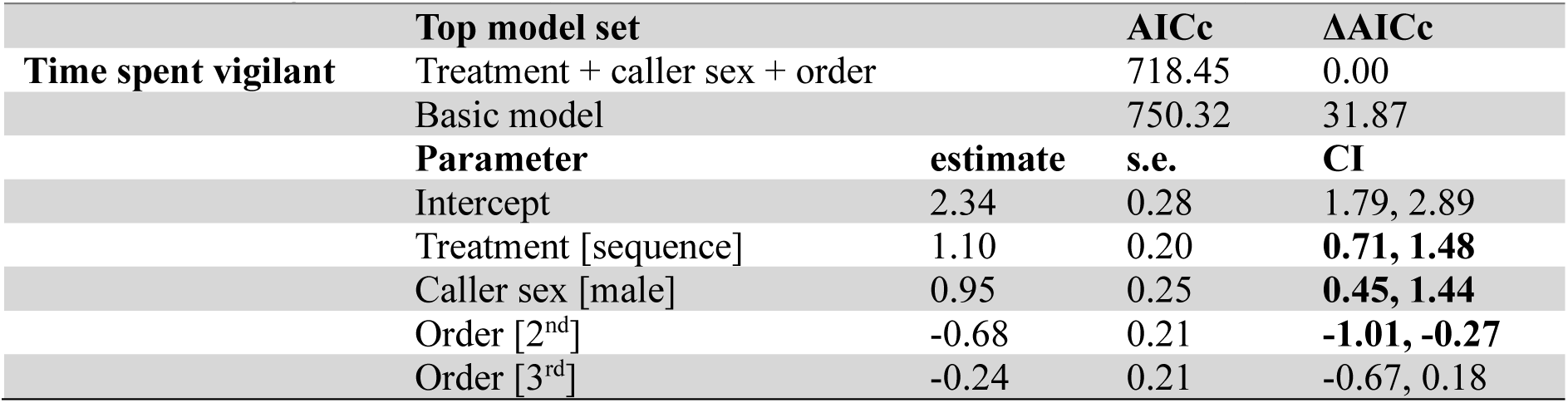
Top model set for time fledglings spent vigilant to a playback experiment (n = 84 trials). Analyses were performed using negative binomial GLMMs with a log link function. The reference levels are alarm for treatment, female for sex and 1^st^ for order. Fledgling ID was included as a random term. Parameter coefficient estimates ± standard error (s.e.) and 95% confidence intervals (CI) are provided for each top model set; see electronic supplementary material for full model sets. Bold CI values indicate significant results (i.e., confidence intervals do not intersect zero).

## Discussion

While ontogenetic research into combinatoriality is gradually emerging (Bortolato et al., 2023; Mason, King & Ridley, 2026; Sigmundson et al., 2025), no study had yet looked at how young first come to understand combinatorial vocalisations. Here, we show magpies exhibit differentiated responses to a discrete call and call sequence very early in development and much sooner than they produce the sequence (Mason, King & Ridley, 2026). In line with our predictions, we showed that fledglings respond with immediate vigilance to both the discrete call and the sequence from the first week of testing (3 weeks post-fledging), supporting the presence of adaptive innate vigilance. Contrary to our predictions however, fledglings also showed differentiated, meaning-derived responses to the sequence—involving prolonged periods of vigilance and increased upward scanning compared to the alarm call—from the first week of testing. Instead of gradual learning of meaning-derived responses, we instead provide evidence that comprehension of a complex vocal sequence emerges rapidly.

The presence of immediate vigilance behaviour following both vocalisations from the start of development is consistent with ontogenetic evidence in other species of early onset vigilance to discrete alarm calls (Hollén & Manser, 2006; Lind & Cresswell, 2005; Wilson & Hare, 2006) and provides novel evidence that this is true of combinatorial vocalisations too. However, the onset of a meaning-derived response to the sequence so early in life is unusual. The majority of research into alarm call comprehension in other social species suggests that while vigilance may be innate or at least emerge early in development, call-specific, meaning-derived behavioural responses generally develop later, following months or even years of associative learning and experience (Hollén & Manser, 2006; León et al., 2023; Seyfarth & Cheney, 1986). Our results suggest either that this period of associative learning occurs far more rapidly in magpies—i.e., in the two weeks between fledging and the start of experimentation—or that there is an innate basis to the response. Given that even the discrete alarm calls of other species typically require some social learning to achieve adult-like comprehension (Hollén & Manser, 2006; León et al., 2023; Seyfarth & Cheney, 1986), it seems unlikely that multi-call sequences—which presumably demand greater cognitive processing—could be innately understood by magpies, especially since these sequences are not innately produced (Mason, King & Ridley, 2026). Given that fledglings could not be tested earlier than week three post-fledge when they began to spend time on the ground, isolated from caregivers, confirming the presence of rapid learning prior to this point would rely on opportunistic natural observations of fledglings in their first two weeks out of the nest responding to both vocalisations. It is of course possible that learning may begin even earlier in the nestling phase (though this would only extend the period of learning opportunity to a still relatively short period of five to six weeks), however, nests are generally high in trees (>15m, personal observation) and semi-concealed by foliage making it unlikely that nestlings have sufficient line of sight to reliably observe adult behavioural responses to natural productions of the sequence. While neither the possibility of rapid learning nor innate comprehension can be ruled out based on the present study, we have provided initial evidence that the comprehension of vocal sequences emerges early in a non-human animal. This lays the groundwork for future research to investigate whether associative learning occurs more rapidly in species capable of producing complex vocalisations, or if there is indeed an innate basis to the comprehension of combinatorial communication.

The fact that fledglings respond differently to the call sequence from as early as three weeks post-fledging—at least seven weeks before any fledgling produced it in the concurrent observational study (Mason, King & Ridley, 2026)—suggests comprehension of sequences, whether learned or innate, precedes the capacity for production. A recent study on the development of vocal sequence production in chimpanzees suggested that vocalisations requiring rapid articulatory change between units within sequences may emerge later in development, in part due to the neuromuscular development required for coordinating production of such vocalisations (Bortolato et al., 2023). While fledglings can produce some of the constituent calls of sequences in isolation from as early as the first week post-fledging (Mason, King & Ridley, 2026), the neuromuscular ‘load’ of combining the calls in quick succession may necessitate a longer period of development. As the same physical maturation is not required for comprehension, this could help explain the earlier onset of comprehension than production of sequences in magpies. Alternatively, delayed production may serve an adaptive function. Premature or inappropriate use of this high-grade threat sequence could have social costs: inexperienced fledglings might erroneously signal an aerial predator, thereby reducing the credibility of the signal and its future effectiveness (Flower et al., 2014; Silvestri et al., 2019; Ydenberg & Lawrence, 1986). This trade-off could help explain the developmental disparity between the onset of comprehension and the onset of production. However, testing whether early production is adaptively suppressed to avoid social penalties is difficult to do directly, for the very reason that fledglings do not produce the sequence during early development. Instead, future work may more fruitfully investigate the presence of cognitive and mechanical constraints on production. Almost all research into neuromuscular development of vocal sequences focusses on birdsong (Adam et al., 2023; Nottebohm, 1971; Riede & Goller, 2010; Suthers et al., 1999) due to the relatively recent discovery of complex non-song combinatoriality in non-human animals. As such, more research is needed to uncover the cognitive and mechanical implications of producing non-song combinatorial vocalisations, and how this may drive differences in the emergence of comprehension and production.

Magpie non-song vocalisations are all composed of the same four vocal segments—*NL, DS, SH* and *LH* (as described above; Walsh et al., 2023). Fledglings produce all four of the vocal segments within either their discrete calls or sequences (Mason, King & Ridley, 2026) and their productions of these segments are acoustically indistinct from those of adults (Mason, 2025). Their presence across all groups and individuals and the lack of acoustic development shows these vocal building blocks are innate (Egnor & Hauser, 2004), with only the way in which they are combined into sequences being socially learned. Paired with the results presented here—showing earlier than expected comprehension of sequences—this raises the question of whether sequences can be decoded by naïve individuals (e.g. young fledglings or out-group adults) based on innately understood composite calls, even if that sequence has not been heard in context previously or is not produced by their social group. For example, a human may have never heard the phrase “as the crow flies” but still be able to decode its probable meaning on first hearing it, due to their understanding of each individual word and the context in which it is said. In this study, only one population-wide sequence was used, due to the availability of high-quality samples from each group, and the need for results to be directly comparable across tested individuals. As such, it was not possible to ascertain whether comprehension of sequences unique to particular groups differs between fledglings from those groups and those from other groups. Future playback experiments using sequences that are unique to specific groups, played to naïve individuals from other groups—especially young fledglings with known consistent group membership—could help determine whether magpies can derive the meaning of novel sequences based solely on their constituent calls.

In addition to finding unprecedentedly early comprehension of these vocalisations, we also found that fledgling reactions were affected by the sex of the caller. Specifically, we found that fledglings consistently spent more time vigilant to playbacks of male callers, regardless of treatment, than those of female callers. Following the completion of data collection for this study, new findings emerged revealing substantial sex differences in the usage of the call sequence used for playback. Walsh (2024) showed that while males reliably produce the sequence in high-threat aerial predator contexts, females often produce it in seemingly nil-threat scenarios (though females reliably used the sequence during group recruitment events and in both contexts an aerial predator or an airborne conspecific intruder may have been present but not visible to observers on the ground). While further observational study of adult usage would be beneficial, the difference in how fledglings respond to male versus female callers could suggest that fledglings are already aware of this sex-specific usage early in development and perceive male callers as having greater ‘signal quality’—i.e., their communicated information is more reliable or at least more consistent with high-level threats (Grüter & Czaczkes, 2019). Adult magpies have similarly been shown to learn to respond less to unreliable callers over time (Silvestri et al., 2019). However, given the alarm call was shown to be used comparably by males and females for general disturbance and group vigilance (Walsh, 2024), we would expect to see an interaction between treatment and caller sex, such that fledglings only show increased vigilance to male productions of the sequence, not the alarm call, but no such interaction was found (Table S.6). One possible alternative explanation is that more frequent aggression towards fledglings from males than females (personal observation) may lead to fledglings being more wary of male vocalisations generally. Again, natural behavioural observations could help to confirm the source of disparity in fledgling responses to male and female callers.

## Conclusion

The results presented here provide the first insight into how combinatorial comprehension emerges during ontogeny. The early onset of meaning-derived responses to combinatorial alarm sequences observed in this study occurs far earlier than the emergence of meaning-derived responses to much simpler alarm calls in other species (Hollén & Manser, 2006). Whether this reflects enhanced cognitive capacity and rapid learning in a communicatively complex species, or points to an innate capacity for decoding complex vocal sequences, remains to be determined. Nevertheless, this study highlights the need to disentangle the respective roles of innate versus learned components in the development of both discrete calls and combinatorial sequences. Understanding this balance is crucial for revealing the cognitive and evolutionary foundations of combinatoriality. By adopting an ontogenetic approach across diverse species, future research may uncover the mechanisms through which combinatorial vocalisations and their meanings emerge across the diverse lineages in which they are found.

