## Supplementary material for "Comprehension precedes production of a complex call sequence"

**Table S.1.** Overview of playbacks done with 9 Western Australian magpie fledglings from 7 groups In Crawley and Guildford, Perth, Western Australia. The alarm treatment was a discrete general disturbance alarm call, and the sequence treatment was a 3-call sequence associated with aerial predators. Location and identity of each group and the identity of the fledgling that was tested at each group are shown. Treatments performed for each (1 fledgling only received the alarm treatment due to the unavailability of sequence exemplars from that group) are shown along with the number of stimuli (i.e. how many individual recordings were used)—these were counterbalanced across weeks tested so the fledglings did not habituate to a single recording. Caller ID shows the identity of the individual whose vocalisations were used to make the stimuli for each fledgling and treatment, along with caller sex. The final column shows which weeks of testing were included in the final dataset—playbacks were conducted every 3 weeks from week 3 to 18 post-fledging (maximum 6 trials per treatment), and some weeks were excluded due to failed playbacks.

| Location | Group | Fledgling ID | Treatment | Number of stimuli | Caller ID | Caller sex | Trials included in final dataset |
|---|---|---|---|---|---|---|---|
| Crawley | BWYb | AMPX | Alarm[1] | 4 | MOTY | M | 4 |
| | | | Sequence[2] | 4 | MOTY | M | 4 |
| | | PYMX | Alarm[1] | 4 | MOTY | M | 4 |
| | | | Sequence[2] | 4 | MOTY | M | 4 |
| | JOG | YXBM | Alarm | 4 | JOG XM | M | 6 |
| | | | Sequence | 2 | JOG XM | M | 6 |
| | MBG | SOLID | Alarm | 3 | VYMY | M | 6 |
| | | | Sequence | 3 | VYMY | M | 6 |
| | SCL | MONKEY | Alarm[3] | 4 | MOBR | M | 6 |
| | | | Sequence | 3 | SCL XF | F | 6 |
| | | MOO | Alarm[3] | 4 | MOBR | M | 5 |
| | | | Sequence | 2 | GEORGE | F | 5 |
| Guildford | NH | NHFL | Alarm | 4 | RMXGMX | F | 5 |
| | | | Sequence | 4 | RMXGMX | F | 6 |
| | PR | PRFL | Alarm | 4 | GRMO | M | 3 |
| | SS | SSFL | Alarm | 3 | SS XM | M | 4 |
| | | | Sequence | 4 | MXXGRY | M | 4 |

Note: Where pairs of treatments for two fledglings are labelled [1, 2,] or [3] the same set of stimuli were used for that treatment for both fledglings.

*Latency to react*

Analysis was conducted using a GLMM with a negative binomial distribution and a log link function. For the top model, no evidence of overdispersion (p = 0.74), zero-inflation (ratioObsSim = 1.91, p = 0.15), or outliers (p =1) was found, and residual normality was confirmed via a Q-Q plot (Kolmogorov-Smirnov test: p = 0.49).

**Table S.2.** Top and full model set for latency to react to playbacks (n = 125 trials). Analyses were performed using negative binomial GLMMs with a log link function. The reference level for treatment is control. Fledgling ID was included as the random term. Parameter coefficient estimates ± standard error (s.e.) and 95% confidence intervals (CI) are provided for each top model set. Bold CI values indicate significant results (i.e., confidence intervals do not intersect zero).

| | **AICc** | **ΔAICc** | **Parameter** | **estimate** | **s.e.** | **CI** |
|---|---|---|---|---|---|---|
| **Top model set** | | | | | | |
| Treatment | 434.41 | 0.00 | Intercept | 3.19 | 0.30 | 2.60, 3.78 |
| | | | Treatment [call] | -4.29 | 0.50 | **-5.27, -3.31** |
| | | | Treatment [sequence] | -3.96 | 0.48 | **-4.90, -3.01** |
| **Full model set** | | | | | | |
| Treatment + proportion spent in contact[1] | 436.47 | 2.06 | | | | |
| Treatment + age[1] | 436.50 | 2.09 | | | | |
| Treatment + average adult association[1] | 436.50 | 2.09 | | | | |
| Treatment + order | 438.32 | 3.91 | | | | |
| Treatment + stimulus | 439.94 | 5.53 | | | | |
| Basic (intercept only) | 495.83 | 61.42 | | | | |

[1] Models had AICc close to being within 2 ΔAICc but were not considered due to main effects having confidence intervals that intersect zero

**Table S.3.** Pairwise comparisons of treatment levels for top model for latency to react conducted using estimated marginal means. Confidence intervals adjusted with a Tukey correction. Estimates and standard errors (s.e.) for respective comparisons are shown. Significant differences are highlighted in bold based on 95% confidence intervals (CI) not intersecting 0. Call is NLDS, sequence is NLDS-SHDS-LHDS and control is the contact call of an Australian wood duck.

| **Response** | **Comparison** | **estimate** | **s.e.** | **CI** |
|---|---|---|---|---|
| Latency to react | Call – sequence | -0.33 | 0.55 | -1.62, 0.96 |
| | Control – call | 4.29 | 0.50 | **3.12, 5.46** |
| | Control – sequence | 3.96 | 0.48 | **2.83, 5.09** |

*Time spent vigilant*

Analysis was conducted using a GLMM with a negative binomial distribution and a log link function. For the top model, no evidence of overdispersion (p = 0.65), zero-inflation (ratioObsSim = 1.77, p = 0.13), or outliers (p =0.84) was found, and residual normality was confirmed via a Q-Q plot (Kolmogorov-Smirnov test: p = 0.88).

**Table S.4.** Top and full model set for time spent vigilant to playbacks (n = 126 trials). Analyses were performed using negative binomial GLMMs with a log link function. The reference level for treatment is control and 1st for order of playback. Fledgling ID was included as the random term. Parameter coefficient estimates ± standard error (s.e.) and 95% confidence intervals (CI) are provided for each top model set. Bold CI values indicate significant results (i.e., confidence intervals do not intersect zero).

| | AICc | ΔAICc | Parameter | estimate | s.e. | CI |
|---|---|---|---|---|---|---|
| **Top model set** | | | | | | |
| Treatment + order [1] | 970.93 | 0.00 | Intercept | 1.91 | 0.20 | 1.52, 2.31 |
| | | | Treatment [call] | 0.15 | 0.19 | **0.79, 1.52** |
| | | | Treatment [sequence] | 2.00 | 0.19 | **1.63, 2.37** |
| | | | Order [2nd] | -0.41 | 0.18 | **-0.77, -0.06** |
| | | | Order [3rd] | -0.21 | 0.19 | -0.57, 0.16 |
| Treatment | 971.52 | 0.59 | Intercept | 1.69 | 0.17 | 1.36, 2.02 |
| | | | Treatment [call] | 1.18 | 0.19 | **0.81, 1.55** |
| | | | Treatment [sequence] | 2.06 | 0.19 | **1.69, 2.44** |
| **Full model set** | | | | | | |
| Treatment + average adult association | 973.41 | 2.48 | | | | |
| Treatment + proportion spent in contact | 973.44 | 2.51 | | | | |
| Treatment + age | 973.62 | 2.69 | | | | |
| Treatment + stimulus ID | 976.18 | 5.25 | | | | |
| Basic (intercept only) | 1046.73 | 75.80 | | | | |

[1] Model not considered because similar AICc to simpler model

**Table S.5.** pairwise comparisons of treatment levels for top model for time spent vigilant conducting using estimated marginal means. Confidence intervals adjusted with a Tukey correction. Estimates and standard errors (s.e.) for respective comparisons are shown. Significant differences are highlighted in bold based on 95% confidence intervals (CI) not intersecting 0. Call is NLDS, sequence is NLDS-SHDS-LHDS and control is the contact call of an Australian wood duck.

| Response | Predictor | Comparison | estimate | s.e. | CI |
|---|---|---|---|---|---|
| Time spent vigilant | Treatment | Call – sequence | -0.88 | 0.18 | **-1.31, -0.46** |
| | | Control – call | -1.18 | 0.19 | **-1.62, -0.74** |
| | | Control – sequence | -2.06 | 0.19 | **-2.51, -1.61** |

*Time spent vigilant—experimental treatments only*

To analyse the effects of caller sex and artificial stimuli (neither of which were relevant to the control treatment, which caused rank deficiency when included), we conducted additional analyses on the experimental treatments only. This only yielded additional results for time spent vigilant–the top predictors for latency to react and proportion spent scanning up remained the same. As such, only the experimental-only model for time spent vigilant is presented. Analysis was conducted using a GLMM with a negative binomial distribution and a log link function. For the top model, no evidence of overdispersion (p = 0.6), zero-inflation (ratioObsSim = 2.34, p = 0.66), or outliers (p =1) was found, and residual normality was confirmed via a Q-Q plot (Kolmogorov-Smirnov test: p = 0.34). All predictors included in the same model had VIF < 2.

**Table S.6.** Top and full model set for time fledglings spent vigilant to two playback treatment levels: a discrete alarm call and a sequence (n = 84 trials). Analyses were performed using negative binomial GLMMs with a log link function. The reference levels are alarm for treatment, female for sex and 1st for order. Fledgling ID was included as a random term. Parameter coefficient estimates ± standard error (s.e.) and 95% confidence intervals (CI) are provided for each top model set. Bold CI values indicate significant results (i.e., confidence intervals do not intersect zero).

| | AICc | ΔAICc | Parameter | estimate | s.e. | CI |
|---|---|---|---|---|---|---|
| **Top model set** | | | | | | |
| Treatment + caller sex + order | 718.45 | 0.00 | Intercept | 2.34 | 0.28 | 1.79, 2.89 |
| | | | Treatment [sequence] | 1.10 | 0.20 | **0.71, 1.48** |
| | | | Caller sex [male] | 0.95 | 0.25 | **0.45, 1.44** |
| | | | Order [2nd] | -0.68 | 0.21 | **-1.01, -0.27** |
| | | | Intercept | 2.34 | 0.28 | 1.79, 2.89 |
| Treatment: caller sex + order [1] | 720.44 | 1.99 | Intercept | 2.54 | 0.39 | 1.77, 3.30 |
| | | | Treatment [sequence] | 0.86 | 0.40 | **0.07, 1.65** |
| | | | Caller sex [male] | 0.75 | 0.37 | **0.02, 1.48** |
| | | | Order [2nd] | -0.70 | 0.21 | **-1.10, -0.30** |
| | | | Order [3rd] | -0.28 | 0.20 | -0.67, 0.11 |
| | | | Treatment [sequence]: caller sex [male] | 0.30 | 0.45 | -0.58, 1.18 |
| **Full model set** | | | | | | |
| Treatment + Age + caller sex + order | 720.89 | 2.44 | | | | |
| Treatment: stimulus type | 721.49 | 3.04 | | | | |
| Treatment + caller sex | 724.21 | 5.76 | | | | |
| Treatment + order | 726.14 | 7.69 | | | | |
| Treatment + Age + caller sex | 726.51 | 8.06 | | | | |
| Treatment | 730.36 | 11.91 | | | | |
| Treatment + average adult association | 732.46 | 14.01 | | | | |
| Treatment + Age | 732.52 | 14.07 | | | | |
| Treatment + proportion spent in contact | 732.55 | 14.10 | | | | |
| Treatment + stimulus ID | 734.86 | 16.41 | | | | |
| Basic (intercept only) | 750.32 | 31.87 | | | | |

[1] Model not considered because of main effects with confidence intervals intersecting zero and similar AICc to simpler model

**Table S.7.** Pairwise comparisons calculated using estimated marginal means of order of playback treatment averaged over caller sex and treatment for the top model for time spent vigilant to two playback treatment levels: a discrete alarm call and a sequence. Confidence intervals adjusted using a Tukey adjustment for multiple comparisons. Estimates and standard errors (s.e.) for respective comparisons are shown. Significant differences are highlighted in bold based on 95% confidence intervals (CI) not intersecting 0.

| Response | Predictor | Comparison | estimate | s.e. | CI |
|---|---|---|---|---|---|
| Time spent vigilant | Trial order | $1^{st}$-$2^{nd}$ | 0.68 | 0.21 | **0.19, 1.17** |
| | | $2^{nd}$-$3^{rd}$ | -0.43 | 0.20 | -0.91, 0.05 |
| | | $1^{st}$-$3^{rd}$ | 0.25 | 0.22 | -0.26, 0.75 |

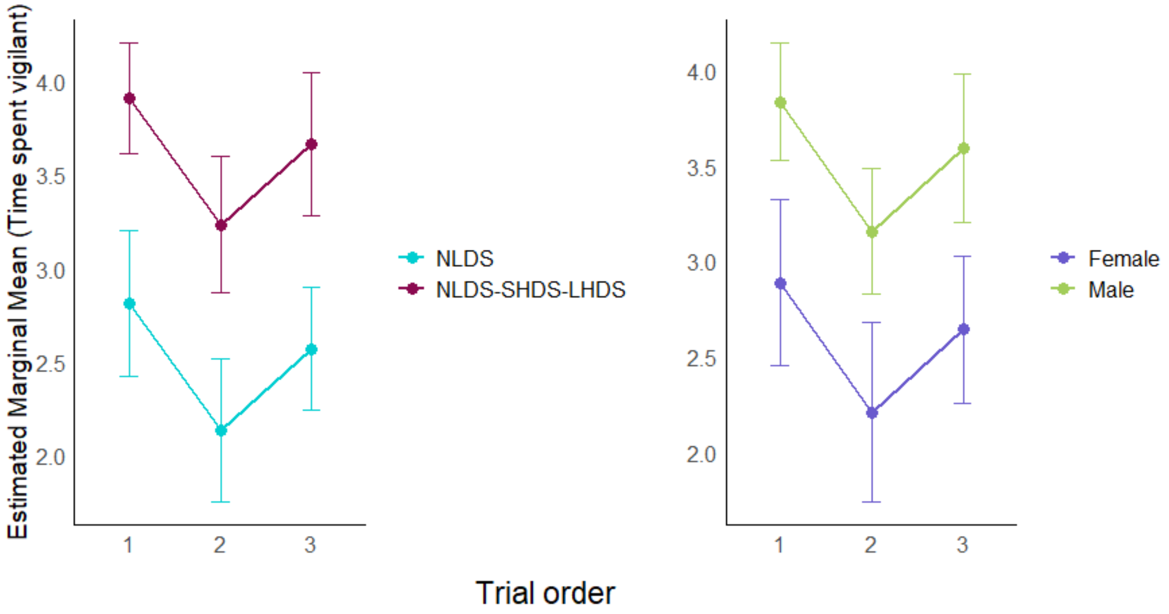


**Figure S.1.** Estimated marginal means of treatment and caller sex when averaged over three levels of trial order (i.e. the position of the treatment in a series of three treatments presented on the day of experimentation). Only two treatment levels are included here: a discrete alarm call (NLDS) and a sequence (NLDS-SHDS-LHDS). Bars illustrate 95% confidence intervals (CI) for estimated marginal means (depicted by dots).

*Scanning upwards*

Analysis was conducted using a GLMM with a binomial GLMM and a logit link function. For the top model, no evidence of overdispersion ($p = 0.70$), or outliers ($p = 0.24$) was found, and residual normality was confirmed via a Q-Q plot (Kolmogorov-Smirnov test: $p = 0.65$). All predictors included in the same model had VIF < 2.

**Table S.8.** Top model set for proportion of vigilance response spent scanning upwards (n = 108 trials). Analysis was performed using a binomial GLMM with a logit link function. The reference level for treatment is control and 1st for order. Fledgling ID was included as a random term. Parameter coefficient estimates ± standard error (s.e.) and 95% confidence intervals (CI) are provided for each top model set. Bold CI values indicate significant results (i.e., confidence intervals do not intersect zero).

| | AICc | ΔAICc | Parameter | estimate | s.e. | CI |
|---|---|---|---|---|---|---|
| **Top model set** | | | | | | |
| Treatment | 518.74 | 0.00 | Intercept | -2.43 | 0.51 | -3.42, -1.43 |
| | | | Treatment [call] | 1.46 | 0.39 | **0.70, 2.22** |
| | | | Treatment [sequence] | 2.61 | 0.37 | **1.89, 3.34** |
| Treatment + order [1] | 518.96 | 0.22 | Intercept | -2.34 | 0.54 | -3.41, -1.28 |
| | | | Treatment [call] | 1.49 | 0.39 | **0.73, 2.25** |
| | | | Treatment [sequence] | 2.59 | 0.37 | **1.86, 3.31** |
| | | | Order [2nd] | 0.04 | 0.16 | -0.28, 0.36 |
| | | | Order [3rd] | -0.30 | 0.17 | -0.64, 0.04 |
| **Full model set** | | | | | | |
| Basic (intercept only) | 657.33 | 138.59 | | | | |

[1] Model not considered because of main effects with confidence intervals intersecting zero and similar AICc to simpler model.

**Table S.9.** Pairwise comparisons calculated using estimated marginal means of proportion of vigilance spent scanning up for each treatment level comparison. Confidence intervals adjusted using a Tukey adjustment for multiple comparisons. Estimates and standard errors (s.e.) for respective comparisons are shown. Significant differences are highlighted in bold based on 95% confidence intervals (CI) not intersecting zero.

| Response | Predictor | Comparison | estimate | s.e. | CI |
|---|---|---|---|---|---|
| Proportion of vigilance scanning up | Treatment | Call – sequence | -1.15 | 0.15 | **-1.50, -0.81** |
| | | Control – call | -1.46 | 0.39 | **-2.37, -0.56** |
| | | Control – sequence | -2.61 | 0.37 | **-3.48, -1.75** |
